# Image-based morphological profiling of autophagy phenotypes in Zika virus infected cells

**DOI:** 10.64898/2025.12.09.693313

**Authors:** Neil Alvin B. Adia, Priya S. Shah

## Abstract

Autophagy is a dynamic intracellular process that is essential in maintaining cellular homeostasis. Its potential as a therapeutic target is exemplified by its dysregulation in many disease states such as Zika virus (ZIKV) infection. ZIKV poses a significant burden to human health and hijacks autophagy to disrupt development. Here, we develop an experimental and computational pipeline to dissect ZIKV hijacking of autophagy in live cells. We build on our previously developed high-throughput scalable image-based profiling approach applied to small molecule perturbation of autophagy. In this study, we expand and modify the image-based profiling pipeline to observe autophagy and virus infection side-by-side using ZIKV encoding a fluorescent reporter. We observe cell and autophagy morphology changes after ZIKV infection and found distinct feature profiles in both infected and uninfected cells treated with ZIKV. We compare ZIKV-infected morphologies to those of cells treated with autophagy-regulating drugs and find ZIKV-induced changes similar to treatment with a combination of autophagy inducer and inhibitor. Using the pipeline, we also train and build a deep learning classifier to identify infected cells using a cellular autophagy reporter without the help of a fluorescent reporter virus. This shows the versatility of image-based profiling in infection-based systems.

## Introduction

Macroautophagy (autophagy) is a dynamic intracellular process that maintains cellular homeostasis through degradation of damaged or unneeded cellular components. It begins when double-membrane structures called phagophores gradually grow around and encapsulate cytoplasmic components. The phagophores eventually become closed double-membrane structures called autophagosomes, which fuse with enzyme-containing lysosomes to form autolysosomes. The lysosomal enzymes degrade components into their basic building blocks, regenerating free amino acids and fatty acids to be used in other processes and pathways [1].

Many diseases dysregulate autophagy, ranging from cancer and neurodegeneration to diseases resulting from virus infection. Autophagy dysregulation during virus infection can happen in different ways. Typically, autophagy is induced as part of the antiviral response by degrading virus particles and components, and some viruses have evolved to inhibit autophagic activity. Moreover, other viruses have gained the ability to manipulate autophagy to their benefit, resulting in a complex relationship involving the interplay between the proviral and antiviral aspects of autophagy [2,3].To illustrate, some flaviviruses can take advantage of autophagy to help throughout their life cycle and can potentially induce or inhibit it depending on how much infection has progressed [4]. One notable example of such a virus is Zika virus (ZIKV), which can cause congenital Zika syndrome, a range of birth defects, in infants. Manipulation of autophagy by ZIKV has been implicated in its fetal pathogenesis [refs]. However, there is still no clear consensus on autophagy dynamics during ZIKV infection, observing both induction and inhibition of the process [5–7]. This may have been limited by the low temporal resolution of the studies, which may not be able to capture all the dynamics of autophagy during infection. Due to this complexity, there is great potential and interest for therapeutic fine-tuning of autophagy during ZIKV infection [8].

Fine-tuning autophagy during virus infection requires careful and precise quantification of both processes. Since both autophagy and virus infection are dynamic and heterogeneous processes, it is important to study them with high temporal frequency and single cell resolution. Live cell experimental systems can provide this resolution. Live cell autophagy monitoring uses microtubule-associated protein 1A/1B light chain 3 (LC3) as a common marker for fluorescent tagging [1,9] while reporter viruses encode an additional protein (usually fluorescent) as a marker. We recently developed a holistic image-based morphological profiling approach to dynamically quantify and characterize autophagy phenotypes in live, single cells after treatment with chemical autophagy modulators [10]. In this study, we modified this approach for the simultaneous quantification and characterization of autophagy phenotypes during ZIKV infection with a fluorescent virus reporter. We found that cells that were exposed to virus showed two distinct modes of morphological changes depending on if they were infected or remained uninfected (bystanders). We compared morphological profiles of bystander and infected cells to those of autophagy drug-treated cells and found some strong correlative relationships with drug treatments, suggesting chemical and viral perturbation of autophagy share common features. We were also able to train and build a deep learning classifier that did not rely on any fluorescent virus image data. While this classifier had modest success (F1 score of 71%), it provides a foundation for viral reporter-free identification of infected cells. In total, this pipeline can be used to visualize and analyze dynamic morphological changes in autophagy that occur during infection, and it shows the potential to be expanded to other combinations of perturbations and cellular proteins.

## Results

### Experimental setup and image-based profiling pipeline

We first adapted our established experimental pipeline to accommodate virus infection. For our original study with autophagy-modulating chemicals, we used a tandem LC3 reporter (pHluorin-mKate2-LC3) to distinguish autophagosomes from autolysosomes. In this new study, we created a single reporter system marking autophagic vesicles (autophagosomes + autolysosomes) using mKate2-LC3. This freed the green channel to track virus infection using a previously developed ZIKV-Venus reporter using the MR766 ZIKV strain [11]. We infected Huh7 human hepatocarcinoma cells expressing mKate2-LC3 with ZIKV-Venus (Figure 1A). This approach can be used with any combination of fluorescent reporter fused to LC3 and a fluorescent reporter virus with different colors. Figure 1B shows illustrative images of cells expressing mKate2-tagged LC3 at 1, 18, and 36 hours post infection with ZIKV-Venus. Images at other timepoints can be found in Figure S1. Since small RNA viruses can evolve quickly, fluorescent reporters are often lost during replication due to their modestly decreased fitness. We therefore tested, through immunofluorescence staining, the co-occurrence of viral antigen and fluorescent reporter. We observed the virus fluorescent reporter was present in all antigen-positive cells, indicating that the reporter is not lost during 48 hours of replication (Figure S2). This was observed in a total of 63 antigen-positive cells visualized in 4 distinct fields on cells plated on a round coverglass.

**Figure 1.**
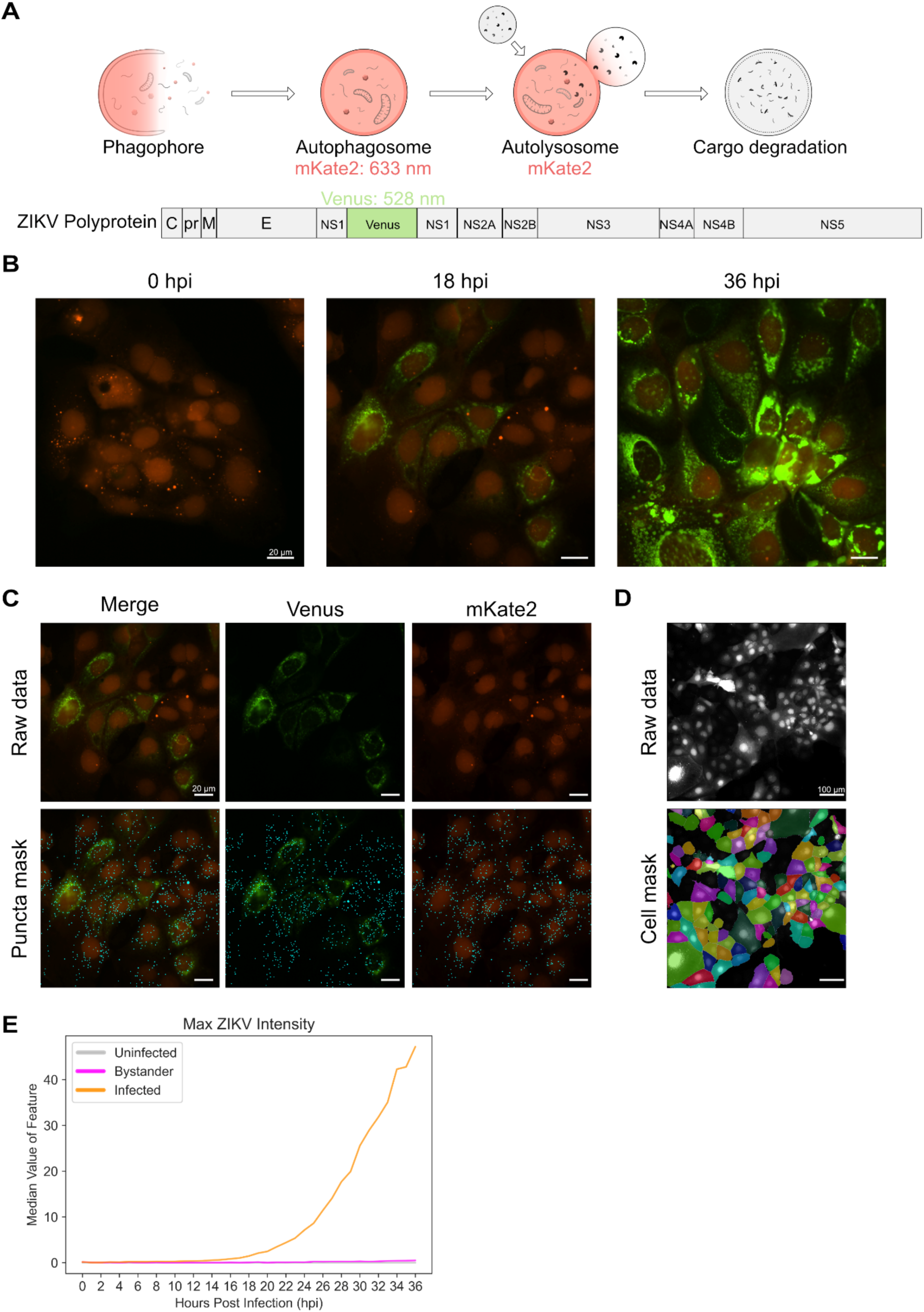
Experimental and analysis pipeline used to study autophagy during ZIKV infection. (A) ZIKV-Venus (GFP channel) was used alongside mKate2-LC3 (TRITC channel) to simultaneously visualize autophagy and infection. (B) Representative images of morphology changes in cells after infection with ZIKV-Venus at different time points. Scale bars represent 20 µm. (C) Representative images of the spot detection algorithm used to detect autophagy vesicles (cyan spots). Scale bars represent 20 µm. (D) Representative image of segmented cell masks used to detect and track single cells. Scale bars represent 100 µm. (E) Line plots showing the median of the maximum GFP intensity in cells for infected, uninfected, and bystander cells.

We next modified the image analysis pipeline developed by Beesabathuni et al. [10] to segment, track, and quantify morphological features of cells (Figure 1C-D). A detailed pipeline highlighting all the major steps is available in Figure S3 and Materials and Methods. Total autophagy vesicle numbers (autophagosomes + autolysosomes) were quantified alongside features belonging to three main categories: structure, intensity, and texture. These features were quantified for the whole cell (Cell), autophagy vesicles (AV), and virus (ZIKV). Infection state for cells was sorted into three categories: cells that were not exposed to virus (uninfected), cells that were exposed to virus but did not get infected (bystander), and cells that were exposed to virus and got infected (infected). Cells that were exposed to virus were filtered by a GFP fluorescence threshold that did not result in false positive cells for the uninfected condition. Figure 1E shows the median value of the mean fluorescence intensity of individual cells at different infection conditions. About 900 features were quantified over the infection time course at a single-cell level.

### Image-based profiling of autophagy features after Zika virus infection

Our previous study involved characterizing morphological changes in cells treated with autophagy inducers and inhibitors [10]. In this study, we infected cells with fluorescent ZIKV with different inoculating ratios known as multiplicity of infection (MOI), which represents the ratio of infecting particles to cells. We used MOI of 0.1, 1, and 3 to represent different regimes of infection in which few, most, or all cells get infected synchronously, respectively. Images were collected every 60 minutes for 36 hours. Images were then processed to extract features for different channels, infection conditions, and time points.

To identify significant changes in features resulting from ZIKV infection, we calculated the Z-score for each morphological feature against the median morphological profile of uninfected cells. This helps standardize the data, not only to help compare features that are scaled differently, but also to reduce training bias in machine learning applications. Features having a median modified Z score >= 0.35 with an adjusted P-value < 0.05 were considered to have changed significantly. These were selected by examining the morphological profiles of mock-infected cells and confirming that a threshold of 0.35 does not classify the features as having changed significantly. This turned out to be a lower threshold than the drug treatments due to the more nuanced nature of perturbations due to virus infection. We show this Z-score analysis at 1, 18, and 36 hours post infection, pooling the infected and bystander cells from different MOIs together (Figure 2A). We annotated the median of the modified Z-score of the number of autophagy vesicles relative to uninfected cells. While the number of autophagy vesicles is not significantly variable at the time points shown, it is worth noting that the median number of autophagy vesicles in bystander cells is lower than in uninfected cells during infection, while infected cells have a higher median number of autophagy vesicles. We also monitored the number of features that varied significantly with time using our Z-score metrics (Figure 2B). Infected cells showed a consistent number of variable features following infection, quickly increasing around the time that GFP signal starts becoming detectable. While ZIKV-related features dominated the changes in infected cells, changes in cell/autophagy vesicle features derived from the mKate2 channel could be identified in infected cells. Bystander cells followed a more erratic pattern of significantly variable features until late time points where they saw a dramatic increase.

**Figure 2.**
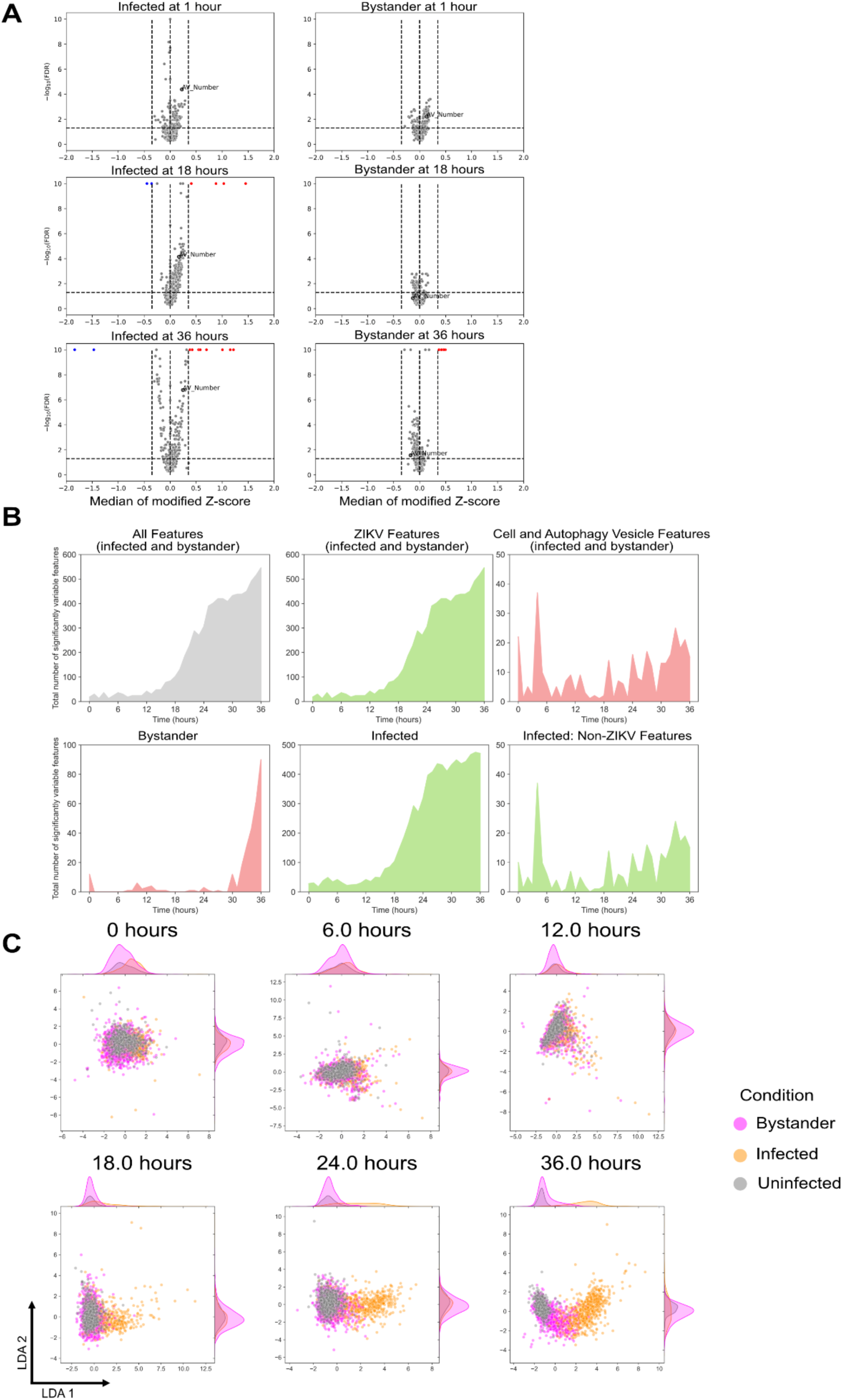
ZIKV infection causes dynamic morphological changes in infected and bystander cells. (A) Volcano plots showing the extent of feature changes for infected and bystander cells at different 1, 18, and 36 hours post infection. (B) Number of morphological features that varied over time for infected and bystander cells, as well as for each channel. (C) LDA plots of infected, bystander, and uninfected cells at different time points.

We visualized the single-cell landscape for infected, uninfected, and bystander cells at different time points using linear discriminant analysis (LDA). Their morphological profiles formed one cluster at the start of infection and eventually segregate into distinct clusters as infection progressed (Figure 2C). Compared to infected cells, bystander cells took longer to split off from the uninfected population, with some remaining similar to uninfected cells in terms of morphology. This may indicate the bystander population includes a combination of cells resistant to infection, cells that are exhibiting paracrine signaling stress responses, and cells that are infected but below our limit of detection.

Our previous analysis with autophagy perturbing drugs showed that integration of temporal data improved separation of cell populations [10]. To improve separation of bystander cells in our study, we visualized the morphological landscape by aggregating all time points on a single LDA plot (Figure 3A, right). Comparing the aggregated analysis with that generated at the single time point of 0 or 36 hours (Figure 3A, middle), aggregated data shows better separation between the three cell states, but more notably between uninfected and bystander cells as shown on the vertical axis. This highlights the strength of time series data when characterizing cellular responses using unsupervised methods.

**Figure 3.**
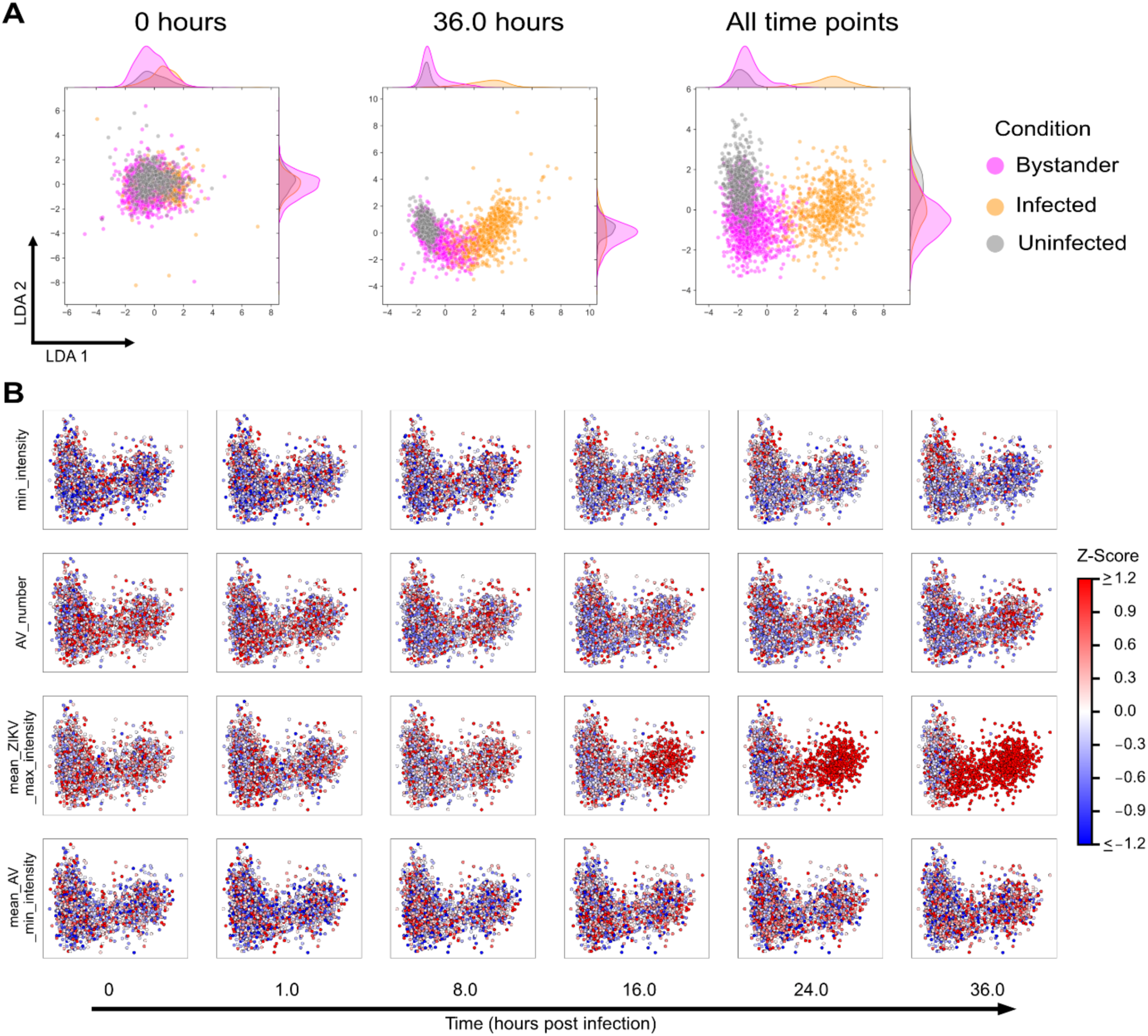
ZIKV infection results in morphological changes for different infection outcomes. (A) LDA plots of infected, bystander, and uninfected cells at 0 hours, 36 hours, and with time-aggregated feature data (all time points). (B) Individual features visualized on the LDA space for different feature categories after infection.

We also visualized individual features on the time-aggregated LDA space (Figure 3B) to track their changes over time for different feature categories (cell, autophagy vesicle dynamics, virus accumulation, and autophagy vesicle fluorescence intensity) that significantly changed over time. While changes in the autophagy vesicle number and the minimum fluorescence intensity of autophagy vesicles were more subtle (decreasing vesicle numbers for bystander cells and overall increasing in brightness for infected and bystander cells), this analysis is also able to show differences in cell responses within the bystander cell population. For instance, infected cells showed an overall decrease in cell fluorescence intensity, while bystander cells showed a more varied change. The bystanders that clustered closer to the infected cell population showed more decreases than those that clustered closer to the uninfected population.

### Image-based profiling shows some similarities between autophagy drug treatment and ZIKV infection

Morphological profile data from well-characterized autophagy modulating chemicals can be a useful and more easily interpretable way to describe cell responses to other less studied perturbations. For ZIKV infection, this was done by sourcing morphological data from Beesabathuni et al. 2025 [10]. The dataset consisted of morphological feature changes after treatment with rapamycin (an autophagy inducer), wortmannin (an autophagy inhibitor), and some combination treatments of both [10]. By making comparisons between the cell and autophagy responses of infected and drug-treated cells, we may gain insight on the types of changes that are being driven by virus infection.

To be able to make comparisons between the two datasets, their morphological feature sets must be reconciled to account for different experimental setups. One major difference was using the green channel to capture autophagosomes versus infected cells. Consequently, GFP features in each set were discarded to reduce inherent artificial separation. The experiments were done with different timelines as well, owing to the different dynamics of the systems. To reconcile this, the time points of drug treatment were linearly scaled [12] so that the final time points of infection would coincide with the final time points of treatment, effectively stretching the dynamics by a constant. Finally, to reconcile autophagy vesicle features, the feature space of drug-treated cells was examined to see whether autophagosome or autolysosome features varied more. In all treatment conditions, more autophagosome features varied compared to autolysosome features. Therefore, autophagy vesicle features from infected cells were compared to autophagosome features in drug-treated cells. The comparison pipeline is visually depicted in Figure S4.

The median morphological profiles of drug-treated and infected cells were dimensionally reduced into the principal component analysis (PCA) space, capturing 90% of the variance within the data. Hierarchical clustering was applied to the data to find the treatments most similar to infection, and Pearson correlation was calculated between the median profiles of the various treatment conditions. Due to the strongly opposing morphological responses between autophagy inducers and inhibitors (Figure S5), infected cell profiles were independently compared with each class of autophagy modulator.

Clustering with rapamycin treatments showed that uninfected and bystander cells mostly showed similarities to untreated cells and cells treated with a very low concentration of rapamycin (Figure 4A), which are expected to produce a relatively negligible change in morphology. While infected cells were shown to cluster with cells treated at moderate concentrations but not high concentrations of rapamycin treatment (Figure 4B), the correlation between their morphologies ranged from weak to moderately anticorrelated (Figure 4C).

**Figure 4.**
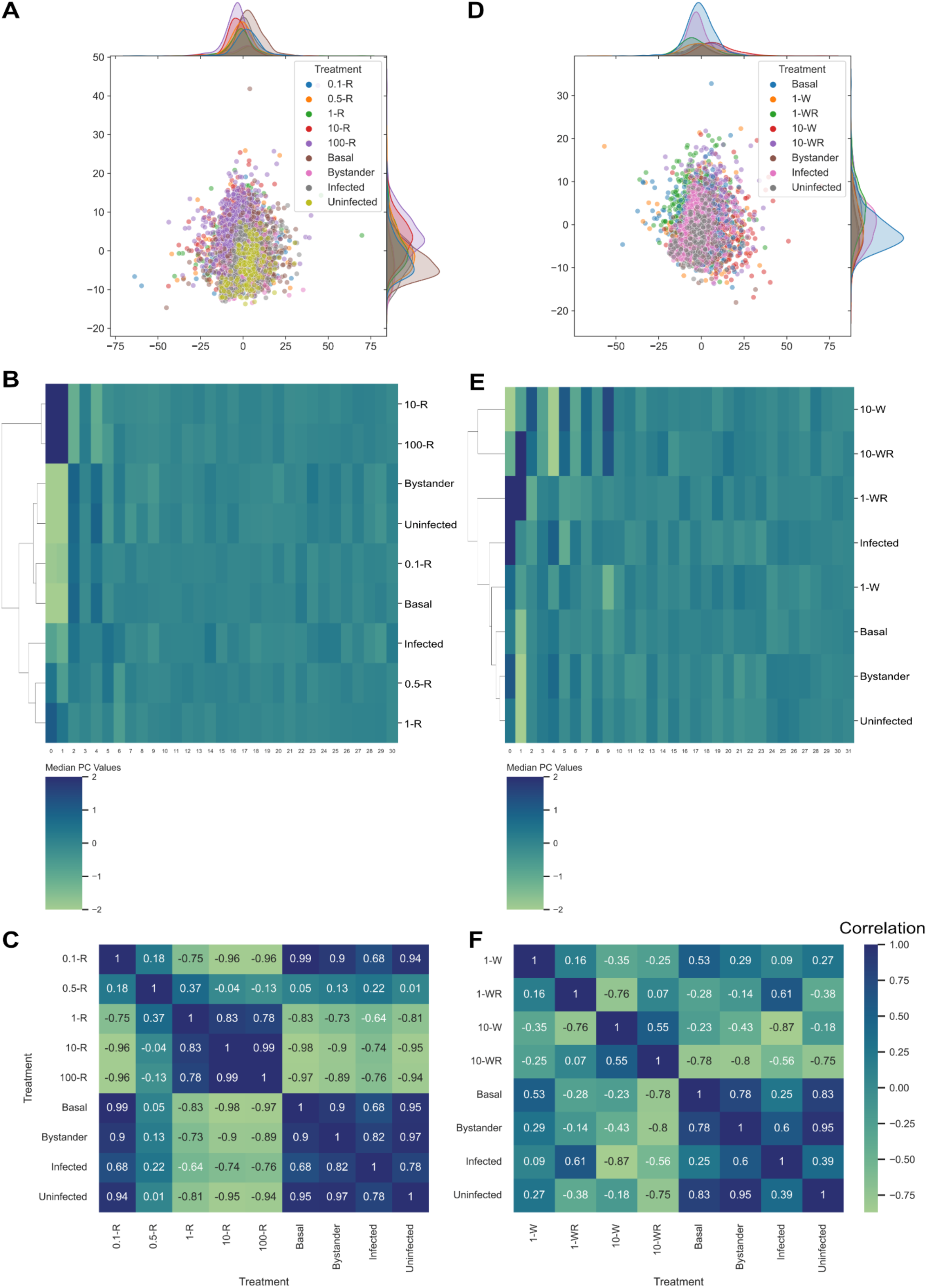
Comparisons of infected, uninfected, and bystander cells to drug-treated treated cells. (A) PCA plot of infected and rapamycin-treated cells. (B) Hierarchical clustering of rapamycin treatment profiles with various infection outcomes. (C) Pearson correlations between rapamycin treatment profiles with various infection outcomes. (D) PCA plot of infected and wortmannin-treated cells. (E) Hierarchical clustering of wortmannin treatment profiles with various infection outcomes. (F) Pearson correlations between wortmannin treatment profiles with various infection outcomes.

Clustering with wortmannin and combination treatments also showed similarities between uninfected, bystander, untreated, and weak wortmannin treatment (Figure 4D). Infected cells did not cluster with strong wortmannin treatment or its combination with rapamycin, but they were more similar to the combination treatment of weak wortmannin and rapamycin (Figure 4E). This combination treatment has the strongest positive correlation with infected cells (Figure 4F), and its autophagy dynamics start with a small decrease in vesicle numbers (wortmannin activity) followed by a sharp increase in vesicle numbers (rapamycin activity) [10].

While this approach makes some assumptions to reconcile disparate datasets, its combination with a semi-automated image analysis pipeline shows an unbiased method for morphological profiles from different perturbations to be reconciled with well-characterized treatments.

### Image-based profiling reveals morphological changes independent of virus reporter

Understanding the differences between infected, uninfected, and bystander cells may shed light on how we can use morphological changes to predict cell fate. Moreover, being able to distinguish between infected and uninfected cells without depending on a fluorescent reporter protein produced by a virus can help in high-throughput diagnostic efforts. It may also open the way to studying cellular morphology for viruses that either do not have an engineered reporter or viruses for which reporter strains are difficult to engineer.

To determine how well autophagic-derived features could be used to identify infected cells, we reanalyzed our data considering only autophagic-derived features from the red channel and discarding virus-derived GFP features. Figure 5A shows an LDA visualization of median morphological profiles for infected, uninfected, and bystander cells with GFP features removed. The horizontal axis shows some slight separation with infected cell profiles from the rest, while the vertical axis shows some separation with uninfected cell profiles from the other cells. The degree of visual overlap is significant between all three conditions, yet the diversity in morphology within infected and bystander cells appears wide. We constructed a random forest model and trained a classifier to identify features that drive the differences between infected and uninfected cells. We optimized design hyperparameters by scanning the parameter space and keeping the configuration that resulted in the highest F1 score, which is a metric that measures a prediction model’s performance in terms of precision and recall. However, classification performance was poor (Figure S6A) which may suggest that individual features may not be enough for classifier training. To account for this possibility, we built and trained a dense neural network to analyze the morphological profiles for each condition (Figure S6B). In a similar manner, we optimized the design hyperparameters for the neural network (see Materials and Methods for more details). After applying 5-fold cross-validation to the data, we then exposed the classifier to a test dataset of at least 100 cells per condition (infected and uninfected) and calculated the F1 score. Training and validation data for the optimized neural network can be found in Figure S6C. The classifier was able to achieve an F1 score of 71%, with a slightly better performance in identifying uninfected cells than infected ones (Figure 5B). This difficulty in classification may be due to the ability of ZIKV to operate stealthily, by evasion from or suppression of cells’ immune capabilities and/or autophagy [4].

**Figure 5.**
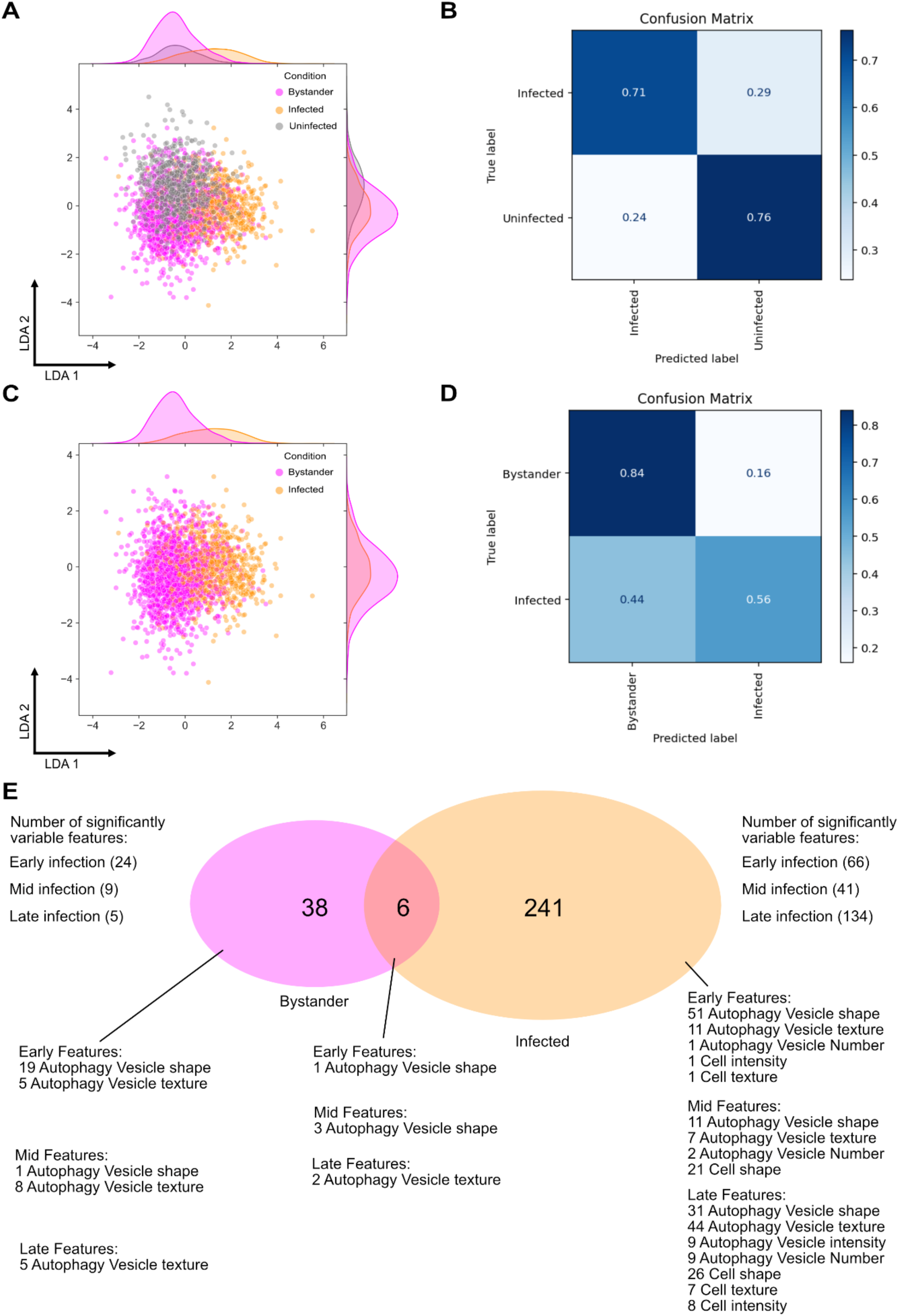
Classification of infected cells using deep learning without the help of a fluorescent virus marker. (A) LDA visualization of cell and autophagy vesicle features for different infection outcomes. (B) Confusion matrix showing the performance of the deep learning classifier in classifying infected vs. uninfected cells without using fluorescent virus data. (C) LDA visualization of cell and autophagy vesicle features for infected and bystander cells. (D) Confusion matrix showing the performance of the deep learning classifier in classifying infected vs. bystander cells without using fluorescent virus data. (E) Cartoon visualization of the features that changed significantly after infection for bystander and infected cells, separated by early (0-11h), mid (12-23) and late (24-36) infection and separated by feature category.

To understand the differences between infected and bystander cells, we visualized their profiles on the LDA space (Figure 5C). We observed key differences in one dimension, but the conditions are largely similar. We also built a dense neural network for classifying between infected and bystander cells with similar model optimization steps, and we found that the model had a higher success rate identifying bystander cells (84%), and more difficulty identifying infected ones (56%) (Figure 5D).

To dissect the key differences in the morphological landscape between the two conditions, we extracted the list of significantly variable features for each condition and split them up into categories to signify early (0-12h), mid (12-24h), and late (24-36h) infection. Figure 5E shows that infected cells have a much larger set of significantly variable features than bystanders. However, it also suggests that the 6 features that they have in common may be playing a more significant role in differentiating between the two.

This pipeline showcases a quantitative, dynamic, and semi-automated approach to studying viruses alongside cellular processes that they hijack. It emphasizes the importance of single-cell measurements that may obscure behavior of bystander cells during infection. It also opens up potential for tracking morphological changes without the need of a fluorescent virus reporter. We anticipate that future efforts expanding on this pipeline, with different viruses or cellular pathways/reporters will add more versatility and predictive power.

## Discussion

Autophagy is dynamic and requires measurement methods that are well-suited to this nature. At the same time, virus infection may consist of dynamic phases. While it is useful to measure on its own, measuring virus infection together with autophagy can contextualize cell and autophagy-specific responses to infection. Furthermore, infection can eventually be paired with drug screening to search for favorable drug candidates at high throughput without expending a large number of resources through destructive sampling or in uncovering fundamental interactions for each drug [13]. Image-based profiling of infection with autophagy also offers a way to examine the heterogeneity of cell phenotypes and responses to infection at a single cell level [10]. This study was a proof-of-concept development of a pipeline to showcase the potential usefulness of image-based profiling to characterize virus infection outcomes.

We expanded systematic side-by-side analysis of infection and autophagy by comparing cell profiles to a morphological screen of autophagy modulation treatments [10]. We found that infected cells produced morphological responses that were vastly different from drug-treated cells that resulted in strong phenotypes or no phenotype at all. Instead, infected cells seemed to follow a more complex morphological pattern that was similar to cells treated with a combination of an autophagy inhibitor and an inducer, finding a moderate correlation between median profiles. This is also reminiscent of other flaviviruses known to modulate autophagy. For example, dengue virus activates autophagy flux early during infection and inhibits it later in infection [14]. Thus, our pipeline refines this understanding of flavivirus perturbation of autophagy with more time and sub-cellular resolution, and with the ability to integrate drug treatment data in a high-throughput manner. We acknowledge that our data integration requires assumptions to reconcile datasets that were generated from processes with different dynamics and different outputs. While future efforts may include generating data with similar experimental setups to avoid these caveats, we emphasize that the approach used here also makes it more likely that existing datasets can be leveraged to their fullest potential.

We tested the pipeline’s potential in not only making predictions of cell infection, but in potentially expanding the utility of this pipeline to study viruses without fluorescent reporters. We built and trained a deep learning classifier with an F1 score of 71% to identify infected vs uninfected cells. Similarly, we trained a classifier to identify infected vs bystander cells, which resulted in a worse performance. While we optimized the hyperparameters to maximize accuracy, we acknowledge the stealthy and heterogenous responses that may emerge from infected and bystander cells in limiting our insight. We examined the effect of relaxing the GFP threshold on the resulting morphological landscape between infected and bystander cells, which further obscured the differences between the two categories when compared without looking at any GFP features (Figure S7).

We also acknowledge that without the use of a classifier similar in principle to random forest models, it will remain a challenge to identify the governing features driving the changes due to infection. However, we are optimistic that efforts in the deep learning space are bridging this interpretability gap through initiatives like Explainable AI [15]. The utility of developing a pipeline without needing a fluorescent reporter virus may also re-enable the use of a dual autophagy reporter, where we can gain insight into the multistep nature of the process and its responses to virus infection, and morphological changes may be more obvious.

We also applied the temporal aggregation of feature data which we previously stated comes with key advantages [10]. We are better able to capture differences in morphology that are otherwise more difficult to discern at individual snapshots of time. This approach also preserves an unbiased and systematic analysis of dynamic processes, especially when critical time points are not necessarily easy to anticipate.

We also acknowledge that the blinded and automated nature of the approach requires thorough manual inspection prior to setting up. These checks involve visual verification of cell and puncta mask generation as well as cell tracking. Additionally, having controls such as uninfected or untreated cells can help ensure that any changes in baseline autophagy by being plated for an extended period can be accounted for. Finally, access to morphological data from well-characterized morphological experiments can also help serve as reference points for expected autophagy behavior.

This pipeline opens up several opportunities when studying viruses and cell responses side by side. It allows for the simultaneous comparison of autophagy responses and changes in virus infection when treating infected cells with autophagy modulators. It may also be combined with pathway-agnostic approaches like Cell Painting [16], to see if any off-target effects may be causing a change in virus infection. Furthermore, protein-protein interaction data for virus and autophagy proteins may be useful in designing experiments to quantify the autophagy response and resulting infection outcomes after the genetic perturbation of certain autophagy proteins [4]. To sum up, this pipeline is an example of how image-based temporal profiling of autophagy can be extended to capture the autophagy response under infection and may serve as a high-throughput method to study viral hijacking of cells.

## Materials and methods

### Cell culture, chemical treatments, viruses, and live cell imaging

Huh7 cells (gift of Dr. Raul Andino) were used for virus infection experiments. HEK 293T cells (gift of Dr. Sam Díaz-Muñoz) were used for packaging lentivirus. Vero cells (ATCC) were used to propagate Zika reporter virus stocks. HeLa cells (accession/identifier) were used in immunofluorescence microscopy. All cell lines were maintained in Dulbecco’s Modified Eagle Medium (DMEM; Thermo Fisher, Gibco) supplemented with 10% fetal bovine serum (FBS; Gibco) inside incubators at 37°C and 5% CO_2_. Cells were washed with Dulbecco’s Phosphate-Buffered Saline (D-PBS; Gibco) and dissociated with 0.05% trypsin-EDTA (Life Technologies). For live-cell imaging, Huh-7 cells were cultured in FluoroBrite DMEM (Gibco) supplemented with 10% FBS and 4 mM of GlutaMAX (Gibco). At least three individual replicates in technical triplicate were performed per condition. Any images with cells that moved out of the focal plane during imaging were manually identified and excluded from analysis using the automated image analysis pipeline.

### Preparation of virus stocks

Fluorescent ZIKV reporter stocks were generated by transfection of pcDNA6.2-MR766-Zika-MR766 3127Intron 2A18VenusNS118 HDVr (gift of Dr. Matthew Evans [11]) into Vero cells, which were monitored for fluorescence and cytopathic effect. Supernatant was harvested and cell debris was removed by centrifugation and then distributed into 500 µL aliquots and frozen at −80°C. Stocks were expanded by propagation in Vero cells and then monitored and harvested similarly. Aliquot titers were determined by plaque assay.

### Virus plaque assay

Vero cells were plated evenly in six-well plates and grown into a monolayer overnight. The following day, virus aliquots were thawed on ice, and 10-fold serial dilutions were performed. Media was removed from the Vero cells, which were then carefully washed with D-PBS. 500 µL of virus dilutions were added to the monolayers and incubated for 1 hour at 37°C with gentle rocking every 15 minutes to ensure even covering. Virus was removed from each well and was replaced by a 3 mL overlay of 1X DMEM mixed with 0.8% methylcellulose (Sigma), 1% FBS, and 1% penicillin/streptomycin (Thomas Scientific) and incubated at 37°C for 4 days. 4% formaldehyde (Thermo Fisher) was then added to the overlay and kept at room temperature for 30 minutes to fix the cells. The formaldehyde-overlay mixture was removed and then the cells were stained with 0.23% crystal violet solution (Thermo Fisher) for 30 minutes. The crystal violet was then removed for plaque counting, and titers were calculated.

### Reporter cell line construction

Lentiviral plasmid FUGW-PK-hLC3 (gift of Isei Tanida; Addgene, 61460; http://n2t.net/addgene:61460; RRID: Addgene_61460) was cut with BamHI and EcoRI. The FUGW backbone sequence was purified and only the mKate2-LC3 sequence of the reporter was amplified by PCR, which was then cloned into FUGW using Gibson assembly, resulting in FUGW-mKate2-LC3. These plasmids were grown in DH5α and Stbl3 cells and purified using MiniPrep (Sigma-Aldritch) or MidiPrep (Macherey-Nagel) kits. Sequences were verified by Sanger sequencing through GeneWiz.

Lentiviral packaging was performed by transfecting 3.5 µg of plasmid into HEK293T with lentiviral packaging plasmids pMDLg/p-RRE (1.8 µg), pCMV-VSV-g (1.25 µg), and pRSV-Rev (1.5 µg), following the calcium phosphate protocol (cite Yu and Schaffer 2006). 36-60 hours post-transfection, lentiviral particles were collected, and cell debris was removed by centrifugation and filtration through a 0.45 µm filter. These stocks were used to transduce Huh7 cells expressing a catalytically dead Cas9 (Huh7-dCas9), generating a cell line we termed Huh7-dCas9-mKate2-LC3. A GFP-encoding lentiviral plasmid was used as a packaging and transduction control. For bulk population sorting, Huh7 cells were harvested and resuspended in PBS supplemented with 1% FBS. Cells positive for mKate2 signal were sorted in a Beckman Coulter Astrios EQ 18-Color cell sorter. A bulk (non-clonal) population of sorted cells was used for all experiments.

### Imaging equipment and settings

Huh7 cells were seeded in 96-well glass-bottom plates with #1.5 cover glass (Cellvis, P96-1.5H-N) and were cultured in FluoroBrite DMEM (Gibco, A1896701) supplemented with 10% FBS and 4 mM GlutaMAX (Gibco, 35050061) approximately 16 hours prior to the start of experiments. Glass bottom plates were first treated with collagen solution (Gibco, A1048301) to increase cell adhesion. Live cell imaging was performed using a Nikon Ti2 Eclipse inverted microscope fitted into an okolab stagetop incubator to maintain the environment at 37°C and 5% CO_2_. Cells were plated at 1.0 × 10^4^ cells per well around 16 hours prior to the start of experiments. NIS-Elements AR software was used to create a semi-automated imaging program to capture images at 2 positions per well at hourly intervals. Images were focused using a Nikon CFI60 Plan Apochromat Lambda 20X Objective Lens, N.A. 0.75, W.D. 1.0 mm, F.O.V. 25mm, DIC, Spring Loaded. Images were captured using an Andor Zyla 4.2 PLUS sCMOS Rolling shutter 53 fps camera. GFP images were captured at a 90 ms exposure and 25% light intensity using a C-FL GFP HC Filter Set, Excitation: 470/40 nm, Emission 525/50 nm, Dichroic Mirror 495 nm. TRITC images were captured at a 350 ms exposure and 30% light intensity. using a C-FL TRITC HC Filter Set, Excitation: 554/23 nm, Emission: 609/54 nm, Dichroic Mirror 565 nm. Illumination was provided by a Lumencore SOLA SE II 365 Light Engine Solid State White-Light Excitation SubSystem. An FPbase configuration of our microscope is available at https://www.fpbase.org/microscope/ESKypKAuDw5fCWiaiedweg/ [17].

### Image processing, cell and puncta mask generation, cell tracking

Images acquired in NIS Elements 5.11.01 are saved as “.nd2” files, which contain all the images and their metadata. The Bright Spot Detection function within NIS Elements’ General Analysis tool was used to generate puncta masks for autophagy vesicles. Rolling Ball background correction was performed using a rolling ball correction radius of 3.90 µm. Parameters used for Bright Spot Detection involved a 1.2 µm typical size and a minimum contrast value of 35. The contrast value accounts for the fluorescence intensity difference between the detected puncta and their surrounding pixels. For larger and unevenly illuminated puncta, the Grow Bright Regions to Intensity. The masked regions for puncta masks were then exported as a separate layer in the “.nd2” file. The general analysis file is available upon request. Examples of spot detection at different time points is shown in Figure S1.

All analysis after puncta mask generation in NIS Elements was done in Python. All supporting code is available on GitHub. Cell masks were generated using the Cellpose algorithm [18], and its human-in-the-loop functionality was used to improve cell segmentation accuracy in an iterative manner [19]. Cell masks were subjected to manual inspection for validation of segmentation accuracy.

Cell tracking was performed using the bTrack algorithm [20] to organize single-cell data temporally. The algorithm takes cell masks as an input and a custom hyperparameter set was used for cell tracks. These hyperparameters are available on GitHub. Cell tracking accuracy was validated by manual inspection for experiments. Feature extraction requires raw images, cell masks, puncta masks, and cell tracks in order to proceed.

### Feature extraction, data preprocessing, standardization, and interpretation

Imaging experiments were set up so that individual images would be named systematically based on the experiment, the replicate, the well, the position in the well, and the channel (GFP or TRITC). This ensured that well positions could be mapped back to their infection condition, and that any replicates done on different days would not have any duplicate labels. This naming system also reduced the complexity for the automation of the image analysis pipeline.

Three sets of morphological features were extracted from cell and puncta masks on raw images: whole cell, virus (ZIKV), and autophagy vesicle (AV). Virus features were extracted from GFP channel images, while whole cell and autophagy vesicle features were extracted from TRITC channel images. The regionprops Python package from skimage.learn was used to extract some features [21]. Haralick texture features and Zernike moments were extracted using the mahotas Python package [22]. All 14 haralick texture features calculated in all directions along with their means and ranges were treated as separate features. 25 Zernike moments were calculated using 0.5*major_axis_length, or half the length of the major axis feature of each cell as the radius. Descriptive statistics for all features for all autophagy vesicles were also calculated using individual vesicle features. These include the mean, median (50^th^ percentile), lower quartile (25^th^ percentile), upper quartile (75^th^ percentile), maximum (max), and minimum (min). A prefix of “ZIKV” was added to virus features, while a prefix of “AV” was added to autophagy vesicle features.

Before preprocessing, all features containing “NaN” values were discarded. To account for batch effects between replicates, every feature extracted for each cell was standardized using the median of mock-infected cells (median(featmock)) then divided by 1.2532 times the mean absolute deviation (MAD(featmock)) of that feature from its respective experiment. The divided value is an approximation of the standard deviation. The resulting data consists of modified Z scores of morphological features.

### Machine learning and dimensionality reduction

PCA analysis was performed using the PCA package from the sklearn.decomposition module to perform clustering of similar treatments. LDA was performed using the sklearn.discriminant_analysis package. Default parameters were used for all dimensionality reduction tasks.

### Random forest classification and feature importance

For random forest classification, features that varied significantly after infection were considered. Features that had a Pearson correlation of 0.75 or higher were removed except for the first one to reduce redundancy. The RandomForestClassifier algorithm from the sklearn.ensemble module was used to make predictions. Classifier accuracy was determined using 5-fold cross-validation. The micro F_1_ score package from sklearn.metrics was used to evaluate classifier performance averaged over 5 iterations. RandomSearchCV was used to search the hyperparameter space for classifier hyperparameters that resulted in the best classification performance. After an optimal configuration was found through random search, GridSearchCV was used on a narrower range to fine-tune parameters. At least 100 cells were used in model testing. The Shapley Additive exPlanations (SHAP) package was used to extract features that were most important for classifier decisions.

### Neural network classification

For classification of images without the virus reporter, a neural network was trained on all features except for those taken from GFP channel images. Initially, all features were taken into consideration, but classification performance was poor. Therefore, features were selected in the same way as random forest classification. The model.predict function from tensorflow.keras was used to make the predictions. To train the model for the classification task, a dense neural network was built, with a 3-layer network used as an initial design basis. The RandomSearch function in the keras_tuner module was used to estimate optimal neural network hyperparameters by generating neural network configurations with random parameter values within specified ranges. 100 random configurations were generated during training, with each configuration being trained twice. The ranges were adjusted iteratively until model performance showed little to no improvement. Neural network hyperparameters included number of neurons per layer, activation function (a function applied to data to amplify positive numbers), regularization (penalizing the model to reduce overfitting), dropout fraction (fraction of neurons to deactivate per iteration to reduce overfitting), number of layers, and learning rate. Early stopping was implemented to reduce the tendency to overfit, and the number of training epochs was also limited to 250. Model accuracy was determined using stratified 5-fold cross-validation. At least 100 cells were used for model testing.

### Hierarchical clustering and Pearson correlation

Both hierarchical clustering and Pearson correlation were based on the combination of all significantly variable features compared to uninfected cells for each condition. The median of these features was considered in the calculation for each treatment. The clustermap function within the seaborn package was used to generate and visualize these plots, using the average method and the euclidean distance metric.

### Statistical analysis

Mann-Whitney U test was used to determine the statistical significance of morphological features of infected cells compared to mock-infected cells. Benjamini-Hochberg Procedure was applied to control the false discovery rate.

## Abbreviations

ZIKV: Zika Virus
LC3: microtubule-associated protein 1A/1B light chain 3
AV: Autophagy Vesicles
GFP: Green Fluorescent Protein
TRITC: Tetramethylrhodamine
LDA: Linear Discriminant Analysis
PCA: Principal Component Analysis

## Author Contributions

NABA conceived the work, executed experiments, developed image and data analysis pipelines, wrote, and edited the manuscript. PSS conceived the work, secured, funding, wrote, and edited the manuscript.

## Data and code availability

Raw feature data (Tables S1-3) and preprocessed data (Table S4) are available on https://figshare.com/articles/dataset/ZIKV_Supplemental/30768023.

Raw image data is available on request due to the excessive size of datasets.

All code for feature extraction, data preprocessing, and image analysis is available on https://github.com/shahlab247/ATG_morphological_profiling/tree/morphology-zikv.

Any additional information that may be needed to reanalyze the data reported in this paper is available on request.

## Acknowledgments

We thank Chase Skawinski, Shruthi Garimella, and Vardhan Peddamallu for their feedback on the manuscript. NIAID R01AI170857 and the W. M. Keck Foundation provided funding for this work.

We also wish to acknowledge the support of the UC Davis Comprehensive Cancer Center Flow Cytometry Shared Resource, supported by the National Cancer Institute of the National Institutes of Health under award number P30CA093373, and S10 OD018223, with technical assistance from Bridget McLaughlin, Jonathan Van Dyke, and Ashley Karajeh.

