## Supplementary Information for "Image-based morphological profiling of autophagy phenotypes in Zika virus infected cells"

#### Supplemental Information

Neil Alvin B. Adia[1], Priya S. Shah[1,2,#]

1 Department of Chemical Engineering, University of California, Davis

2 Department of Microbiology and Molecular Genetics, University of California, Davis

Key words: autophagy, image-based profiling, Zika virus infection, machine learning

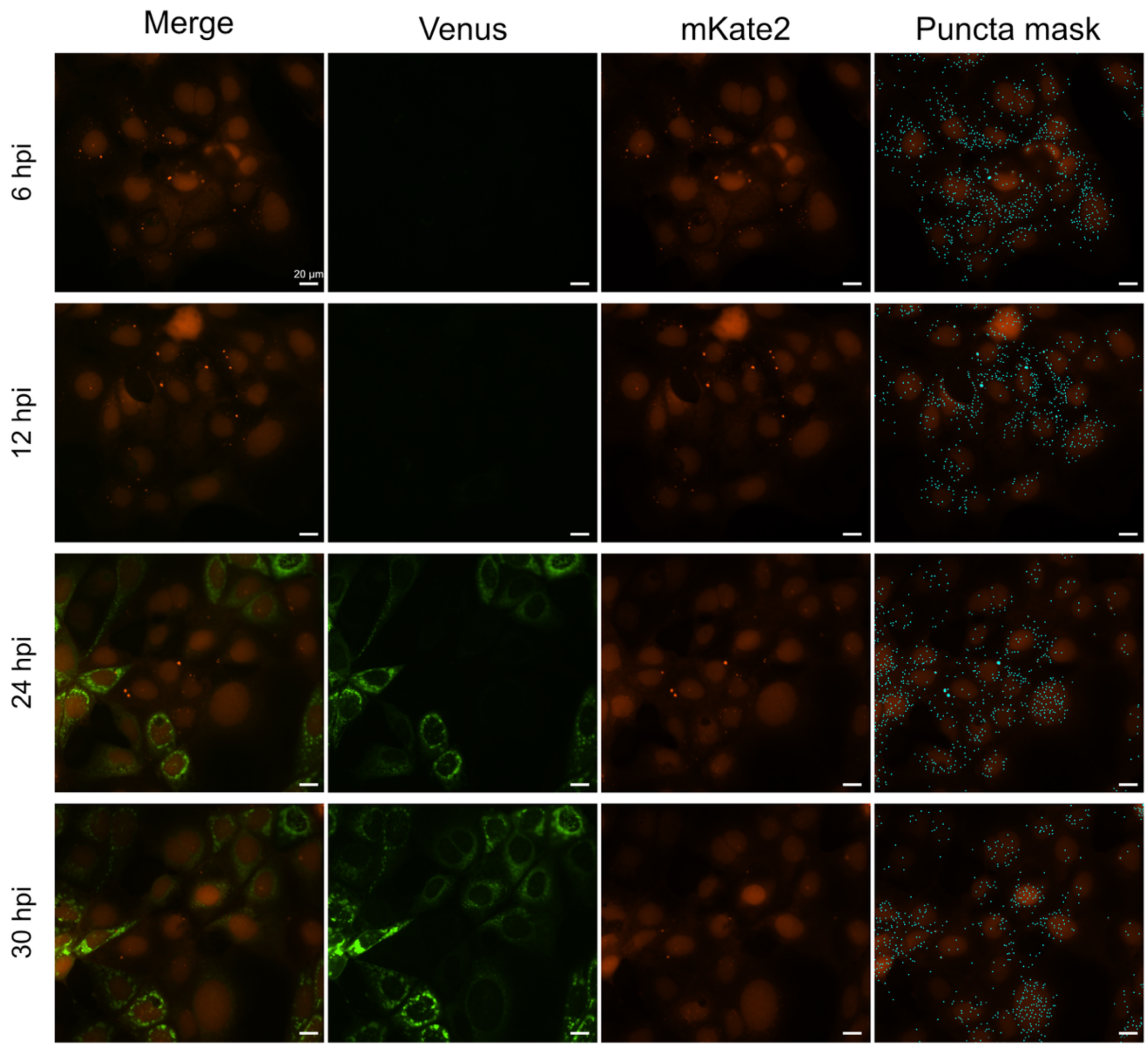


Figure S1 Representative images of morphology changes and puncta mask generation after ZIKV infection.

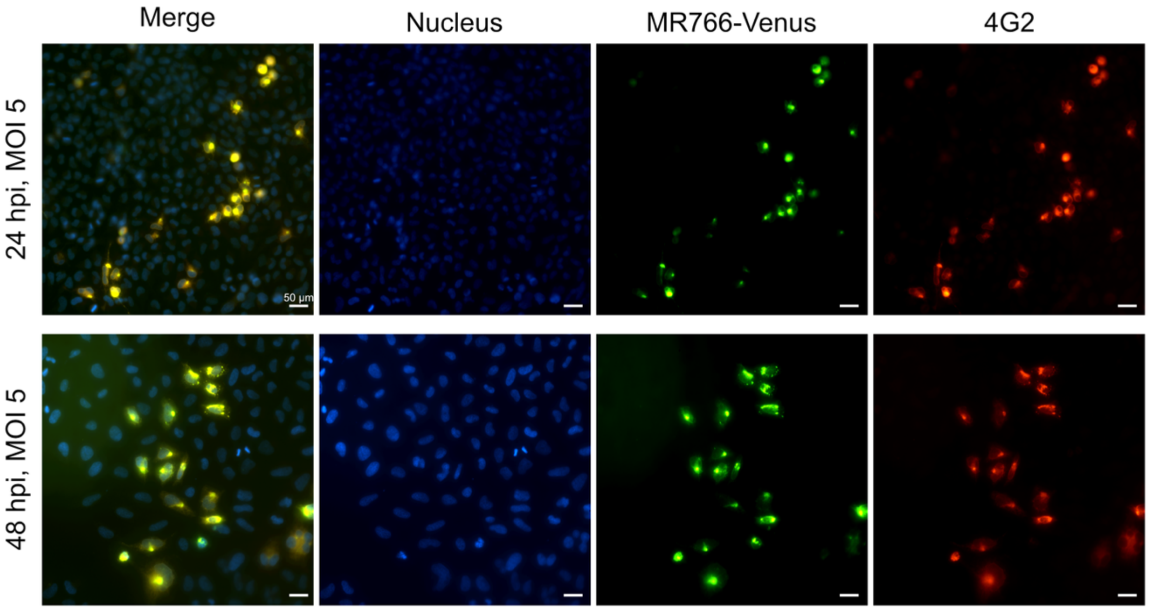


Figure S2 Immunofluorescence staining of ZIKV at 24 and 48 hours post infection to show that Venus is not lost during infection.

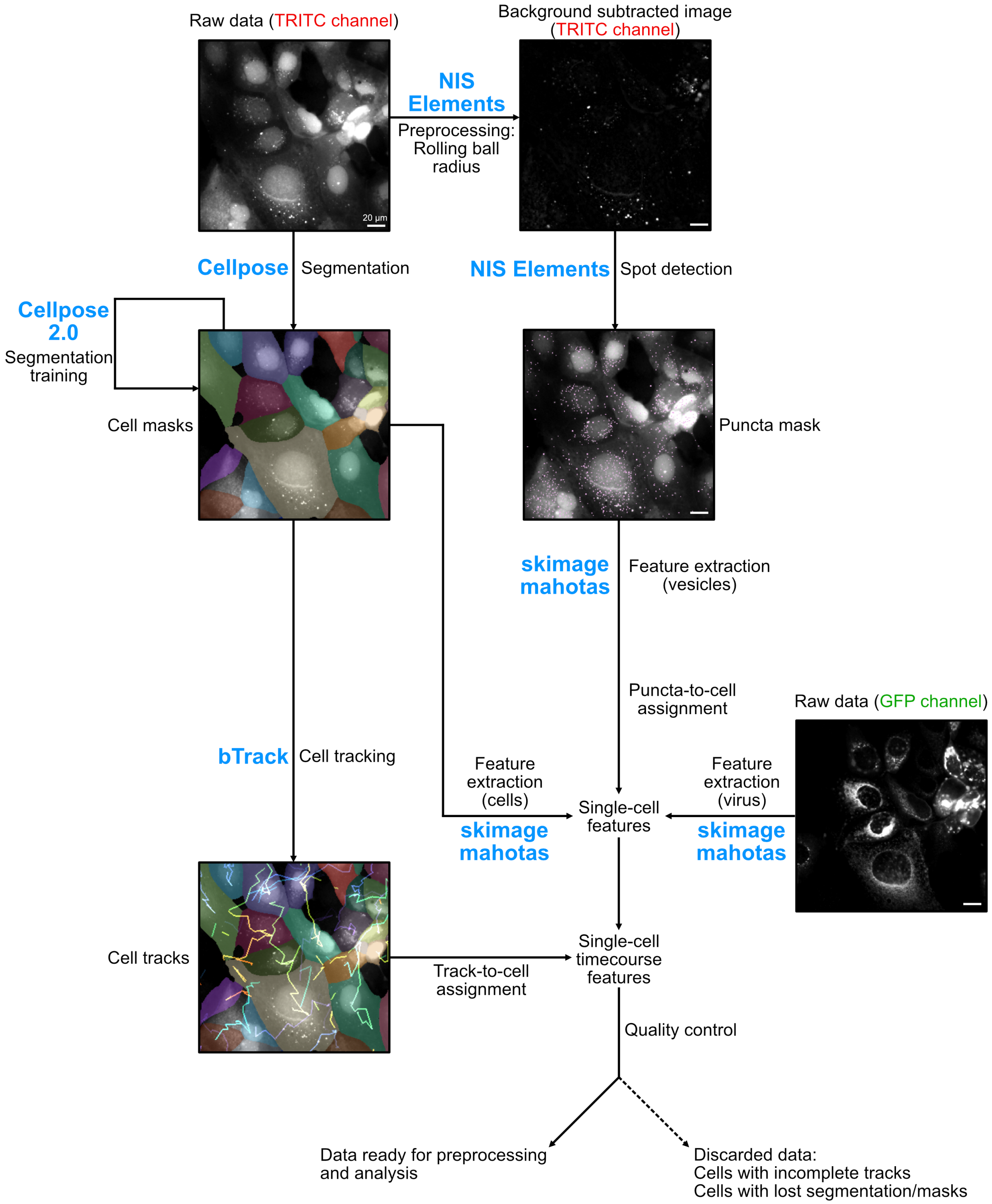


Figure S3 Image analysis pipeline for virus-infected cells.

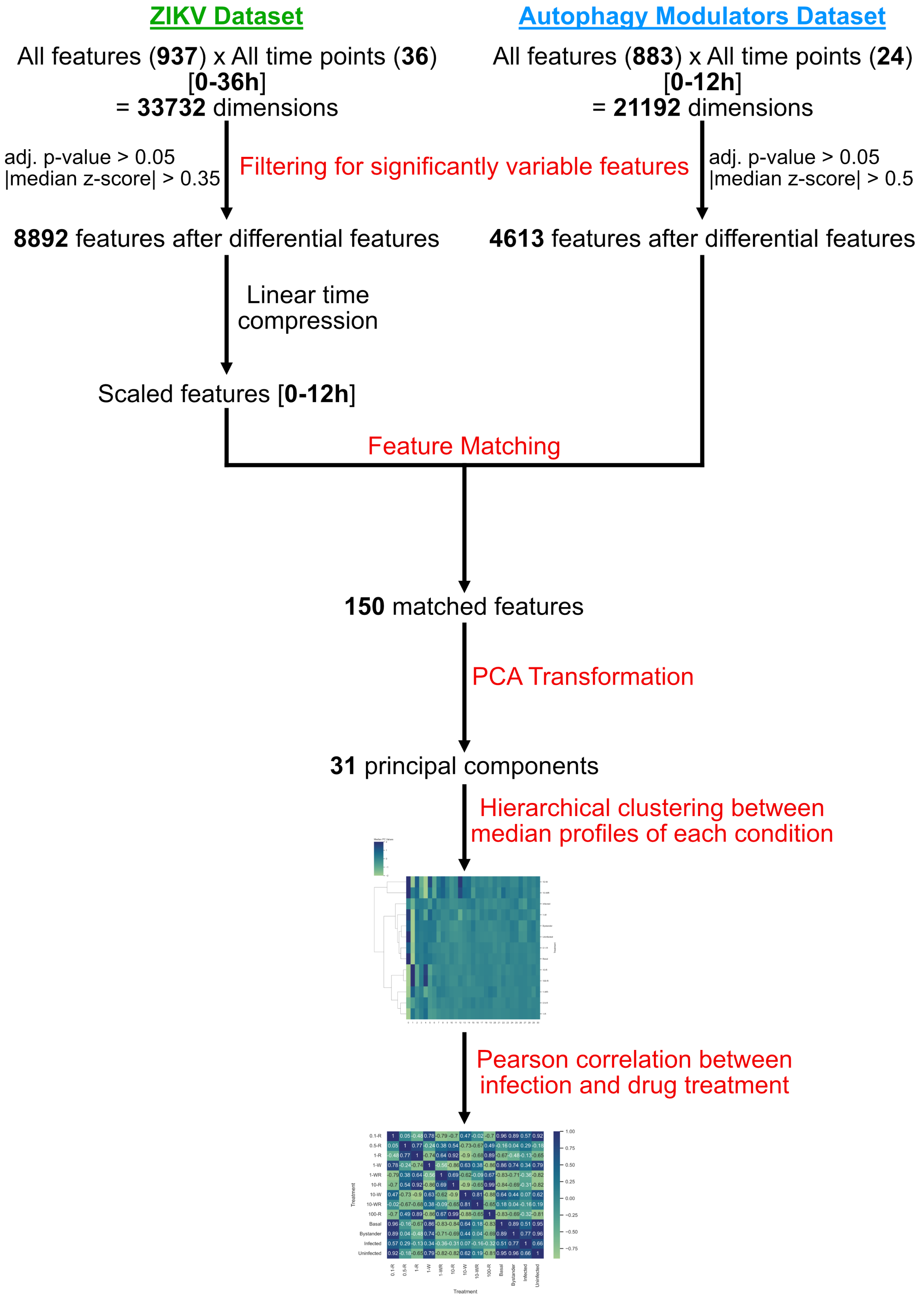
ZIKV Dataset
Autophagy Modulators Dataset
All features (937) x All time points (36) [0-36h] = 33732 dimensions
All features (883) x All time points (24) [0-12h] = 21192 dimensions
adj. p-value > 0.05
|median z-score| > 0.35
Filtering for significantly variable features
adj. p-value > 0.05
|median z-score| > 0.5
8892 features after differential features
4613 features after differential features
Linear time compression
Scaled features [0-12h]
Feature Matching
150 matched features
PCA Transformation
31 principal components
Hierarchical clustering between median profiles of each condition
Pearson correlation between infection and drug treatment

Figure S4 Pipeline for reconciling disparate data sets of infected and drug-treated cells.

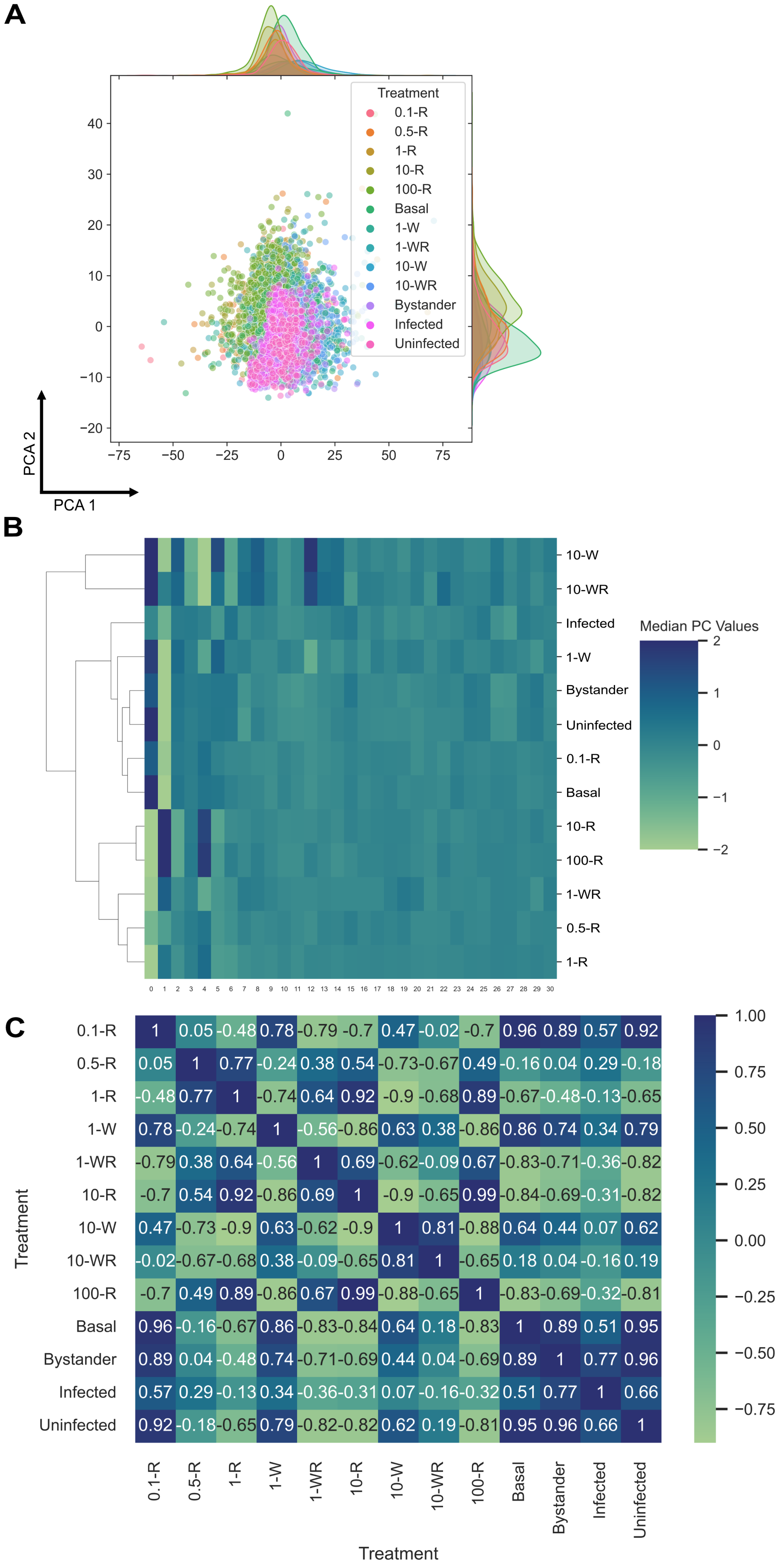
A
Treatment
0.1-R
0.5-R
1-R
10-R
100-R
Basal
1-W
1-WR
10-W
10-WR
Bystander
Infected
Uninfected
PCA 2
PCA 1
B
Median PC Values
10-W
10-WR
Infected
1-W
Bystander
Uninfected
0.1-R
Basal
10-R
100-R
1-WR
0.5-R
1-R
C
Treatment
0.1-R 1 0.05 -0.48 0.78 -0.79 -0.7 0.47 -0.02 -0.7 0.96 0.89 0.57 0.92
0.5-R 0.05 1 0.77 -0.24 0.38 0.54 -0.73 -0.67 0.49 -0.16 0.04 0.29 -0.18
1-R -0.48 0.77 1 -0.74 0.64 0.92 -0.9 -0.68 0.89 -0.67 -0.48 -0.13 -0.65
1-W 0.78 -0.24 -0.74 1 -0.56 -0.86 0.63 0.38 -0.86 0.86 0.74 0.34 0.79
1-WR -0.79 0.38 0.64 -0.56 1 0.69 -0.62 -0.09 0.67 -0.83 -0.71 -0.36 -0.82
10-R -0.7 0.54 0.92 -0.86 0.69 1 -0.9 -0.65 0.99 -0.84 -0.69 -0.31 -0.82
10-W 0.47 -0.73 -0.9 0.63 -0.62 -0.9 1 0.81 -0.88 0.64 0.44 0.07 0.62
10-WR -0.02 -0.67 -0.68 0.38 -0.09 -0.65 0.81 1 -0.65 0.18 0.04 -0.16 0.19
100-R -0.7 0.49 0.89 -0.86 0.67 0.99 -0.88 -0.65 1 -0.83 -0.69 -0.32 -0.81
Basal 0.96 -0.16 -0.67 0.86 -0.83 -0.84 0.64 0.18 -0.83 1 0.89 0.51 0.95
Bystander 0.89 0.04 -0.48 0.74 -0.71 -0.69 0.44 0.04 -0.69 0.89 1 0.77 0.96
Infected 0.57 0.29 -0.13 0.34 -0.36 -0.31 0.07 -0.16 -0.32 0.51 0.77 1 0.66
Uninfected 0.92 -0.18 -0.65 0.79 -0.82 -0.82 0.62 0.19 -0.81 0.95 0.96 0.66 1
0.1-R 0.5-R 1-R 1-W 1-WR 10-R 10-W 10-WR 100-R Basal Bystander Infected Uninfected
Treatment

Figure S5 Comparisons of infected, uninfected, and bystander cells to drug-treated treated cells. (A) PCA plot of infected and drug-treated cells. (B) Hierarchical clustering of drug treatment profiles with various infection outcomes. (C) Pearson correlations between drug treatment profiles with various infection outcomes.

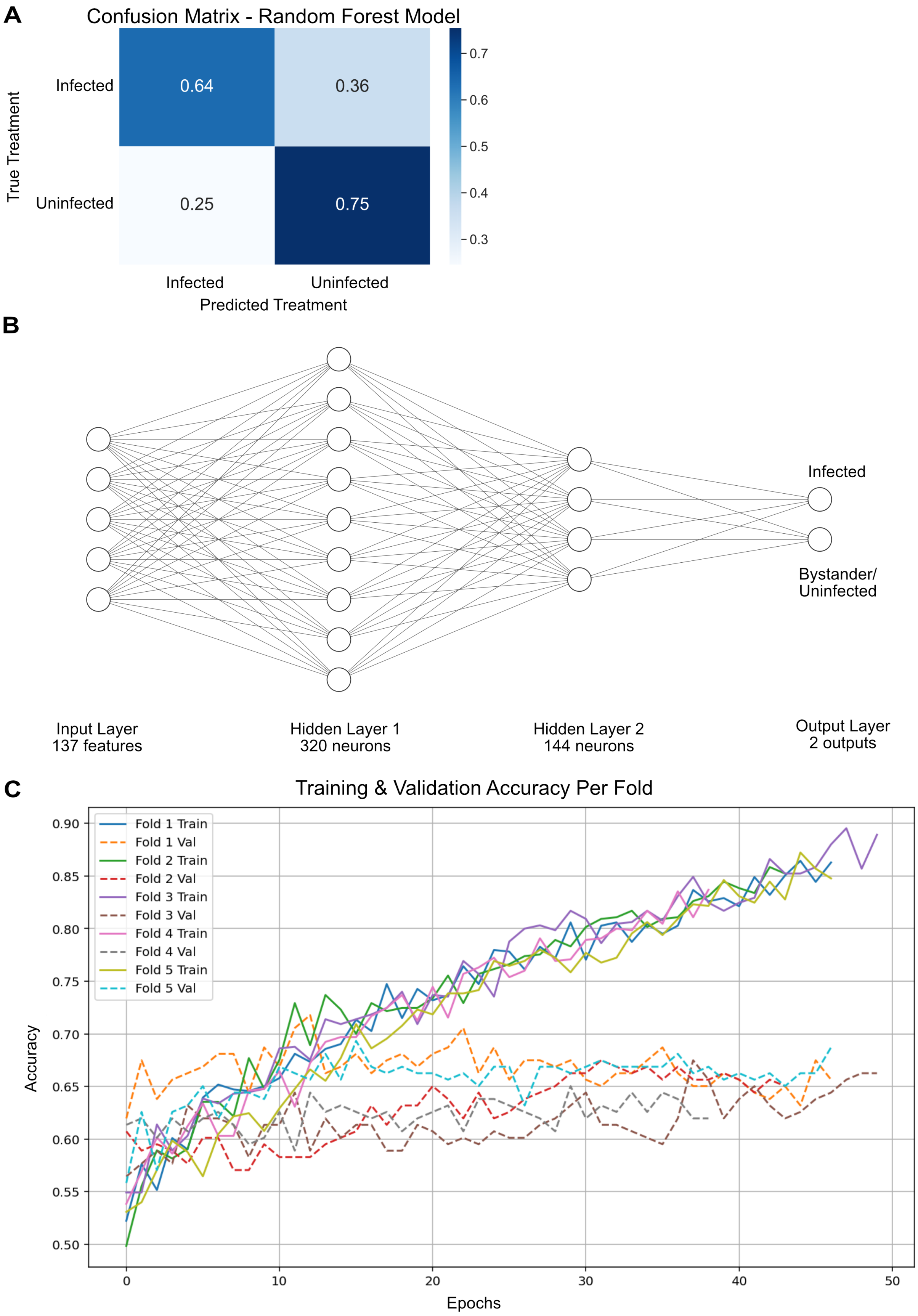
A
Confusion Matrix - Random Forest Model
True Treatment
Infected
Uninfected
0.64
0.36
0.25
0.75
Infected
Uninfected
Predicted Treatment
0.7
0.6
0.5
0.4
0.3
B
Infected
Bystander/
Uninfected
Input Layer
137 features
Hidden Layer 1
320 neurons
Hidden Layer 2
144 neurons
Output Layer
2 outputs
C
Training & Validation Accuracy Per Fold
Fold 1 Train
Fold 1 Val
Fold 2 Train
Fold 2 Val
Fold 3 Train
Fold 3 Val
Fold 4 Train
Fold 4 Val
Fold 5 Train
Fold 5 Val
Accuracy
Epochs
0.90
0.85
0.80
0.75
0.70
0.65
0.60
0.55
0.50
0
10
20
30
40
50

Figure S6 Design and performance of a deep learning classifier. (A) Performance of an optimized Random Forest classifier to identify infected cells without the help of a fluorescent virus marker. (B) Dense neural network architecture for the deep learning classifier. (C) Training and validation accuracy and performance using K-fold cross validation.

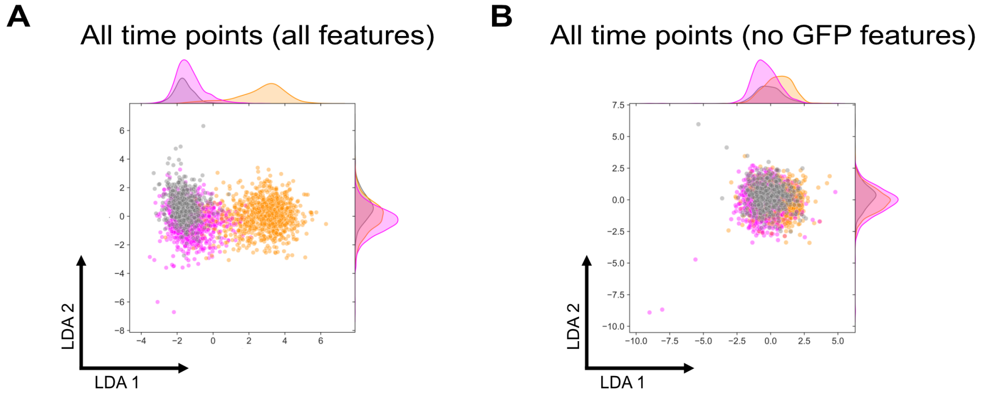


Figure S7. Effect of relaxing the GFP threshold on the morphological landscape of infected and bystander cells. (A) LDA plot representing the complete morphological profiles of infected, uninfected, and bystander cells at all time points after a relaxed GFP threshold. (B) LDA plot representing morphological profiles of infected, uninfected, and bystander cells at all time points without any virus-derived features after a relaxed GFP threshold.

Supp Figure 1

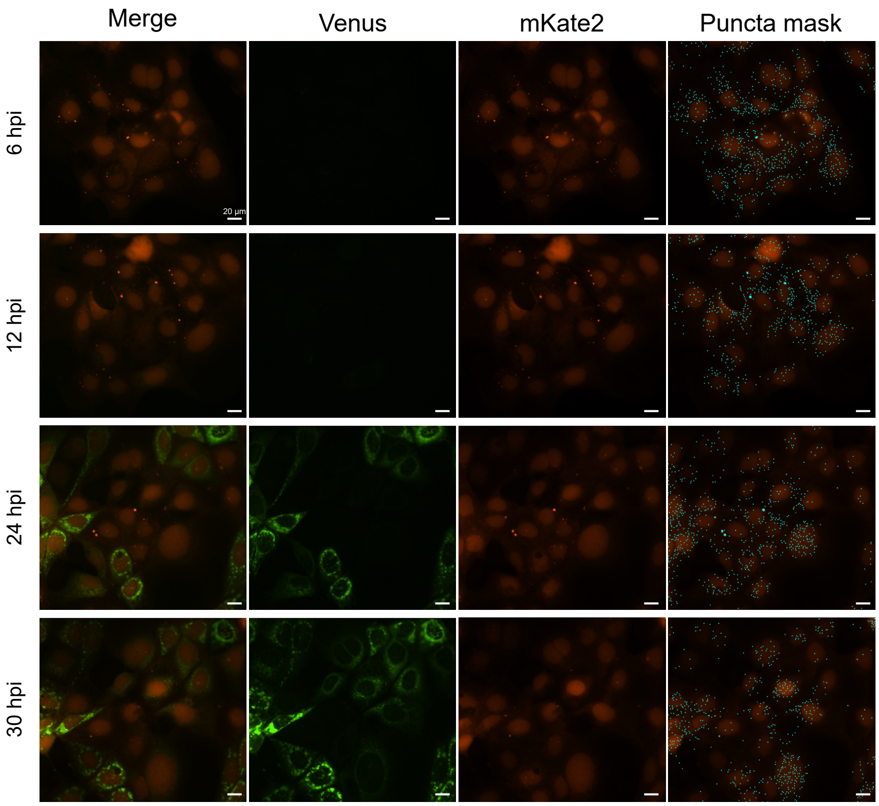

Supp Figure 2

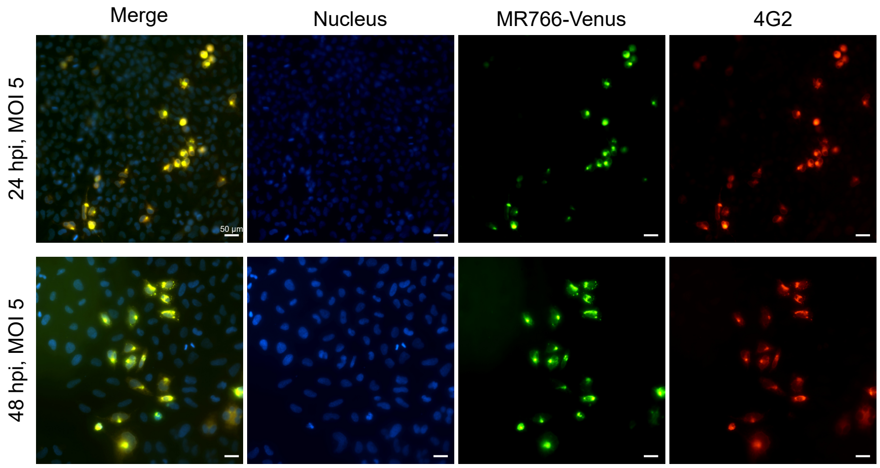

Supp Figure 3

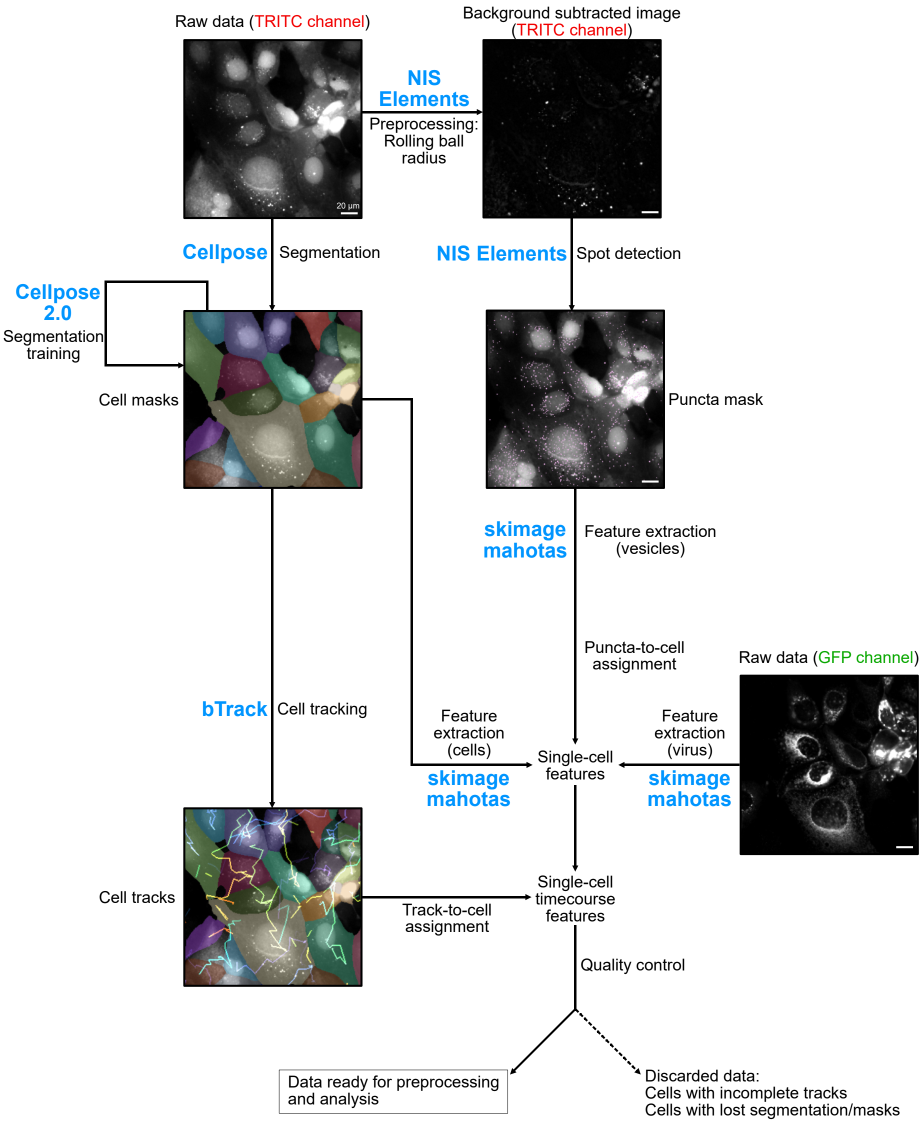

Supp Figure 4

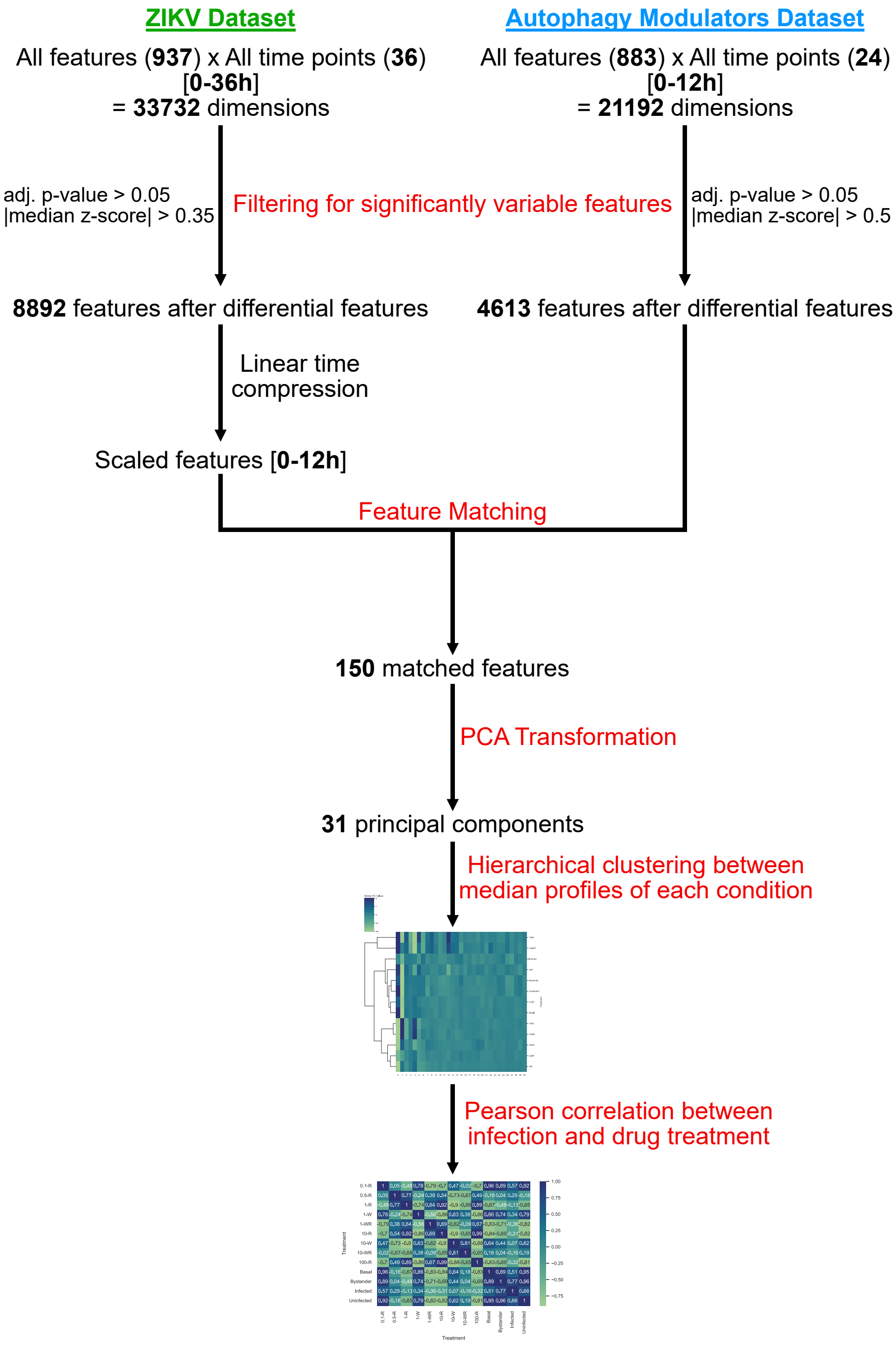

Supp Figure 5

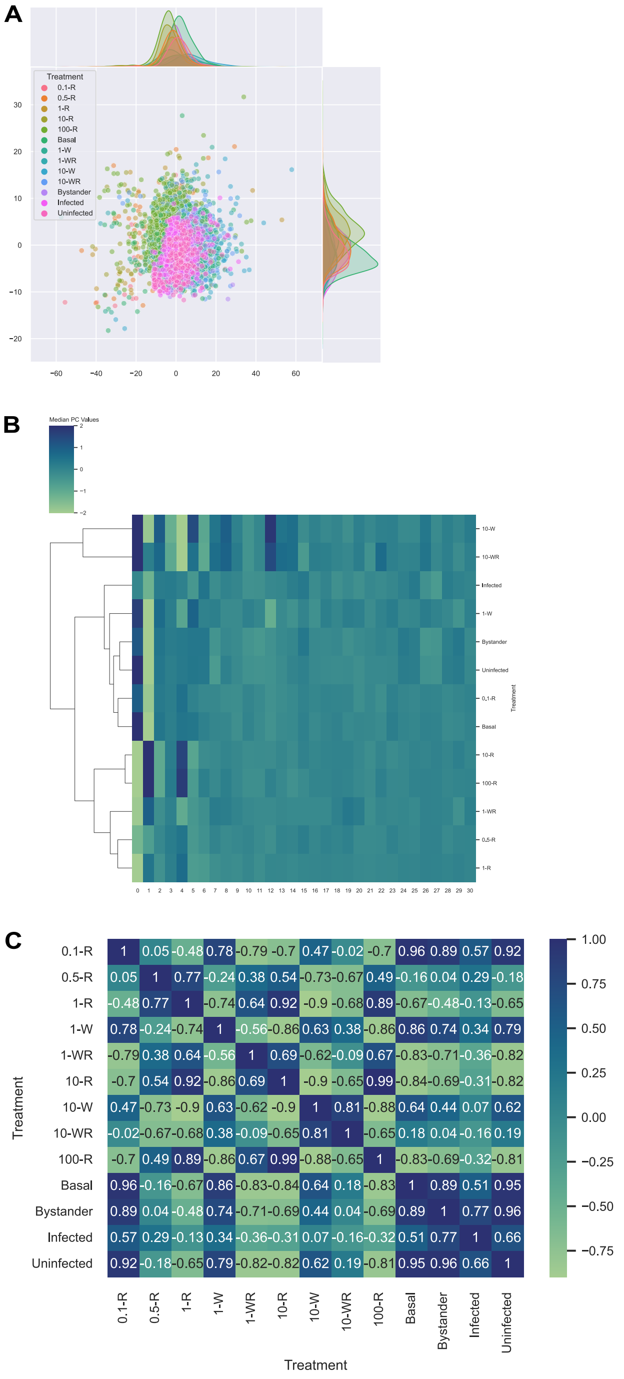

Supp Figure 6

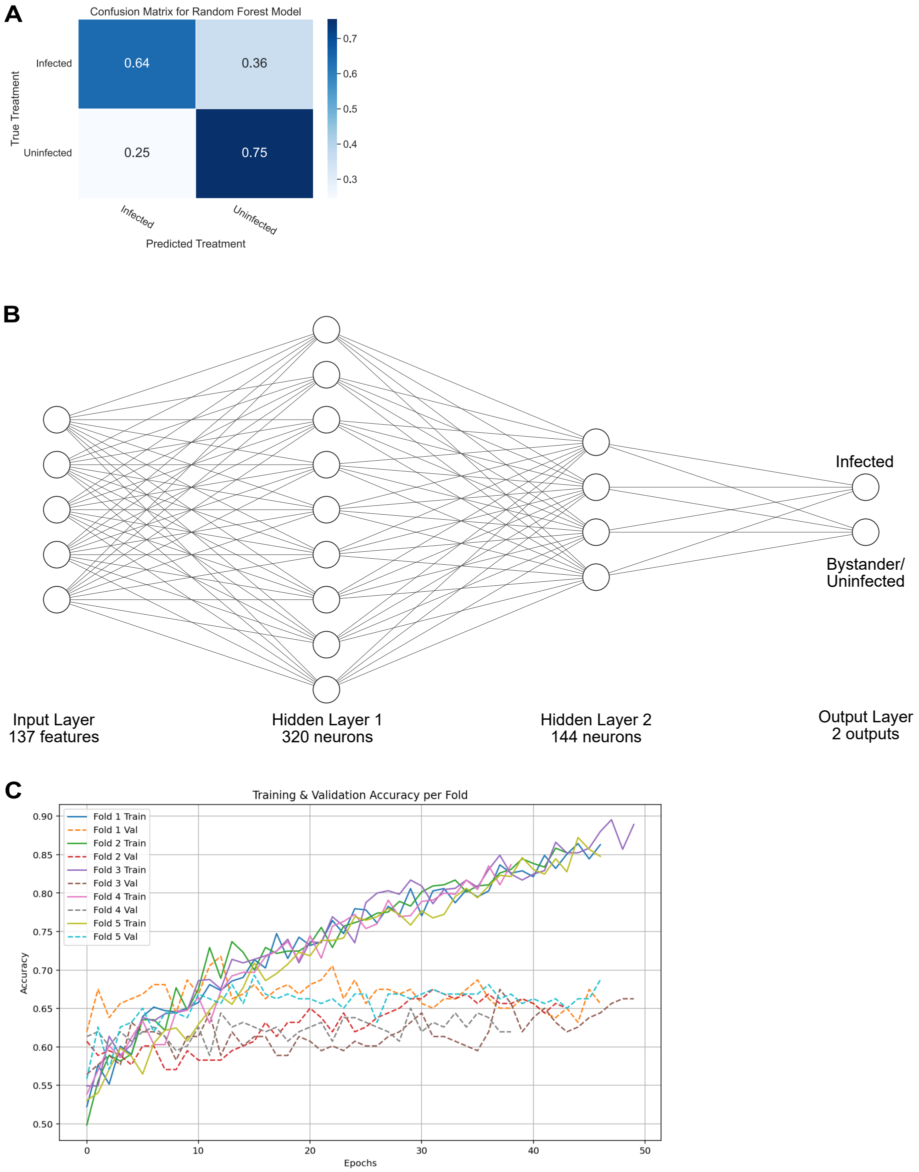

Supp Figure 7

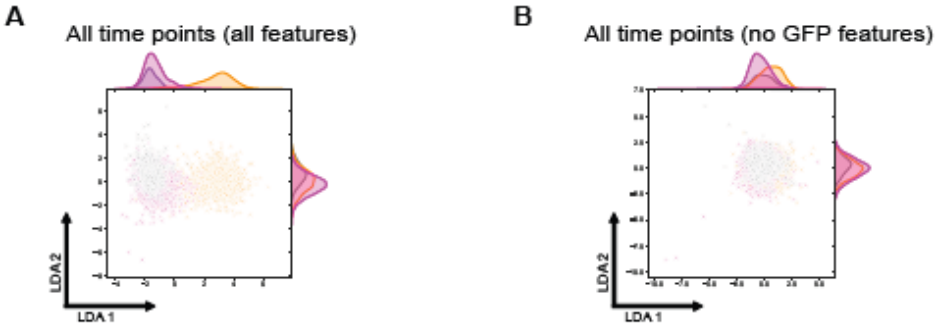
